# Characterizing a Lethal CAG-ACE2 Transgenic Mouse Model for SARS-CoV-2 infection with Using Cas9-Enhanced Nanopore Sequencing

**DOI:** 10.1101/2024.05.30.596396

**Authors:** Alexander Smirnov, Artem Nurislamov, Galina Koncevaya, Irina Serova, Evelyn Kabirova, Eduard Chuyko, Ekaterina Maltceva, Maxim Savoskin, Daniil Zadorozhny, Victor A. Svyatchenko, Elena V. Protopopova, Oleg S. Taranov, Stanislav S. Legostaev, Valery B. Loktev, Oleg Serov, Nariman Battulin

## Abstract

The SARS-CoV-2 pandemic has underscored the necessity for functional transgenic animal models for testing. Mouse lines with overexpression of the human receptor ACE2 serve as the primary animal model to study COVID-19 infection. Overexpression of *ACE2* under a strong ubiquitous promoter facilitates convenient and sensitive testing of COVID-19 pathology. We performed pronuclear microinjections using a 5 kb CAG-ACE2 linear transgene construct and identified three founder lines with 140, 72, and 73 copies, respectively. Two of these lines were further analyzed for *ACE2* expression profiles and sensitivity to SARS-CoV-2 infection. Both lines expressed *ACE2* in all organs analyzed. Embryonic fibroblast cell lines derived from transgenic embryos demonstrated severe cytopathic effects following infection, even at low doses of SARS-CoV-2 (0,1-1.0 TCID_50_). Infected mice from the two lines began to show COVID-19 symptoms three days post-infection and succumbed between days 4 and 7. Histological examination of lung tissues from terminally ill mice revealed severe pathological alterations. To further characterize the integration site in one of the lines, we applied Nanopore sequencing combined with Cas9 enrichment to examine the internal transgene concatemer structure. Oxford Nanopore sequencing (ONT) is becoming the gold standard for transgene insert characterization, but it is relatively inefficient without targeted region enrichment. We digested genomic DNA with Cas9 and gRNA against the *ACE2* transgene to create ends suitable for ONT adapter ligation. ONT data analysis revealed that most of the transgene copies were arranged in a head-to-tail configuration, with palindromic junctions being rare. We also detected occasional plasmid backbone fragments within the concatemer, likely co-purified during transgene gel extraction, which is a common occurrence in pronuclear microinjections.

## Introduction

Transgenic animals are widely used to model various diseases. For coronavirus infections, such as SARS-CoV and SARS-CoV-2, the presence of receptor proteins on animal cells is crucial for virus internalization. Specifically, angiotensin-converting enzyme 2 (ACE2) interacts with the receptor-binding domain of the virus spike protein, facilitating endocytosis and internalization (Kuba et al. 2010). Many strains of transgenic mice with human ACE2 have been created to model COVID-19 (reviewed in (Battulin and Serov 2022; Clever and Volz 2023).

The most popular model, CK18-hACE2, uses tissue-specific regulatory elements, such as the 5′-promoter and the first intron of the human *cytokeratin-18* (*CK18*) gene, the viral translational enhancer and the 3’-part of the *CK18*, to ensure transgene activity in epithelial cells of the respiratory tract and lungs (McCray et al. 2007). Recent studies have shown that CK18-hACE2 mice infected with SARS-CoV-2 exhibit symptoms similar to human COVID-19 (Yinda et al. 2021). Additionally, the HFH4 promoter, specific for ciliated lung epithelial cells, has been used effectively in *ACE2* genetic constructs (Menachery et al. 2016). These transgenic mice have also been exploited to test antiviral therapies against COVID-19 (Jiang et al. 2020).

As an alternative, the mouse *ACE2* promoter has been effective for creating tissue-specific models (Yang et al. 2007). These lines exhibited the expected human *ACE2* expression profile (lungs, heart, kidneys, small intestine) and virus replication without lethality, unlike most other mouse models (Bao et al. 2020). Precision genetic modification using CRISPR/Cas9 technology has been reported by two groups, leading to site-specific knock-ins at the mouse *ACE2* locus (Sun et al. 2020; Liu et al. 2021). These humanized mouse models show typical COVID-19 infection patterns but vary in symptoms due to different co-integrated regulatory elements at the *ACE2* locus. These models also could survive COVID-19 infection.

An alternative strategy involves using constitutive ubiquitous promoters rather than tissue-specific ones. While this approach might not accurately reproduce COVID-19 disease progression due to a wider range of affected tissues, it offers interesting research options. Transgene expression variability can lead to unique infection responses. These models are also convenient for testing vaccines and protective antibodies due to the severe disease course. Several ACE2 mouse lines using the CAG promoter have demonstrated its effectiveness (Tseng et al. 2007; Dolskiy et al. 2021; Bruter et al. 2021).

In this study, we describe the generation of a murine model susceptible to SARS-CoV-2 through the widespread expression of human *ACE2* driven by the CAG promoter. Infection of these transgenic mice leads to rapid death.

Additionally, we discuss the applicability of the Nanopore enrichment method, based on a single cut inside the transgene, rather than two cuts at the flanks like in Nanopore Cas9-targeted sequencing (nCATs) (Gilpatrick et al. 2020).

## Results

### Creation of the CAG-ACE2 mouse line

To create mice susceptible to SARS-CoV-2, we selected a classical approach to overexpress the human ACE2 receptor through which the virus enters cells. The particular *ACE2* coding sequence was taken from the hACE2 plasmid (Addgene #1786), that was successfully tested before in a multitude of studies (Li et al. 2003). Since the goal was also to ensure a severe course of the disease, we decided to use a strong constitutive ubiquitous CAG promoter to drive expression. The diagram of the created genetic construct is shown in Fig. 1A. For microinjections, we used a linear fragment cut from the plasmid containing chimeric CAG promoter cDNA of human *ACE2* and a polyadenylation site from rabbit beta globin gene.

**Figure 1.**
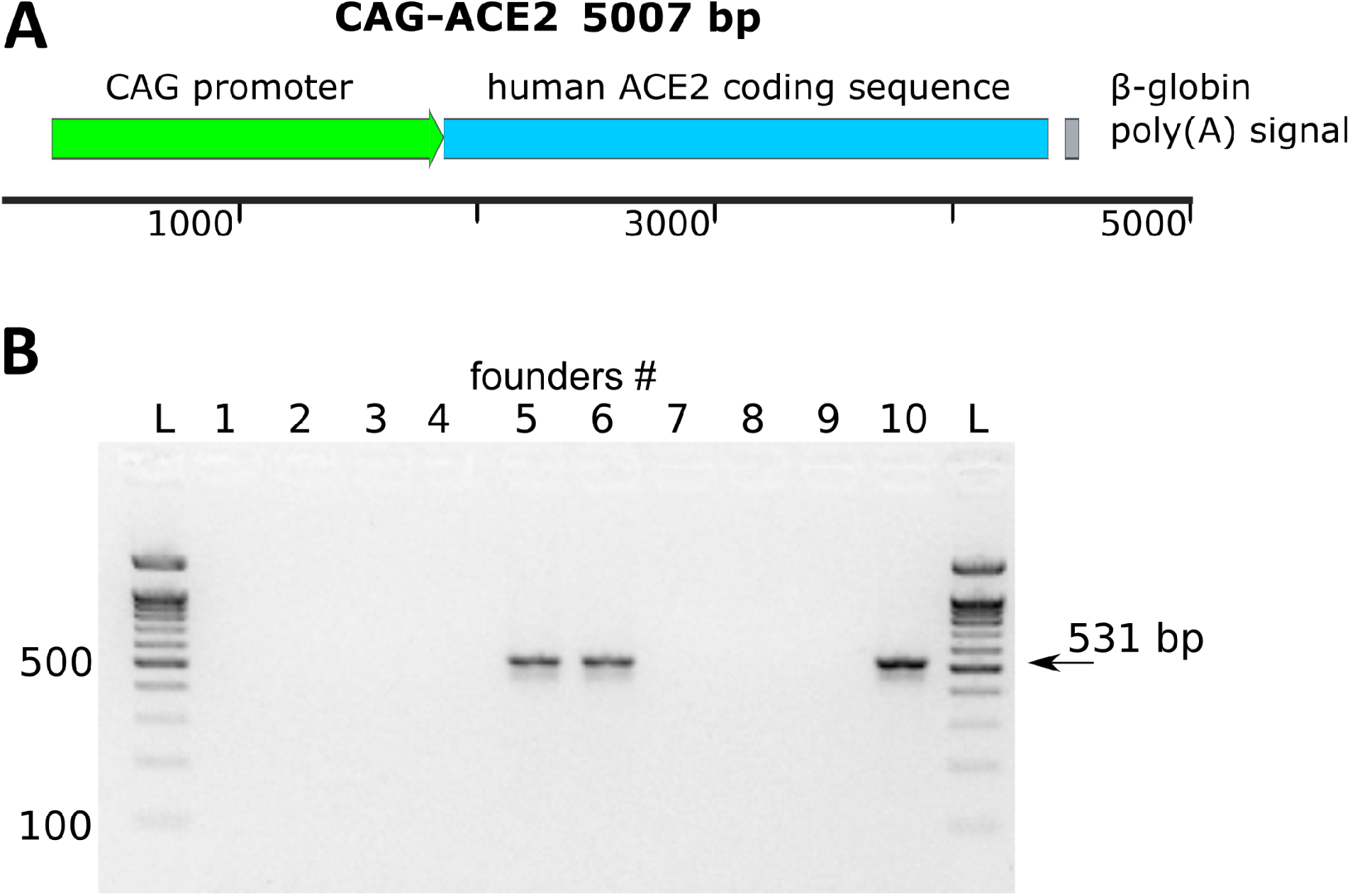
Generation of transgenic animals expressing ACE2. (A) ACE2 transgene structure. Scale bar denotes DNA length (bp). (B) Identification of 3 transgenic animals among 10 F0 offspring. The expected size of the PCR product is 531 bp. L - DNA ladder (100 bp).

We performed 244 pronuclear microinjections using the design described above. 173 embryos successfully underwent this procedure and were transplanted into 11 pseudopregnant females, of which 4 successfully gave birth to 10 offspring. PCR analysis showed that 3 of 10 pups were transgenic (Fig. 1B). Three transgenic lines (#5, #6, #10) were obtained from the founders by crossing with the parental line C57/BL6, but further we will concentrate on the description of the two of them (#5 and #6). We did not observe any significant differences in the properties of these lines in the following experiments.

### The CAG promoter mediates ubiquitous ACE2 activity in transgenic mouse tissues

We analyzed transgene activity in the tissues of F1 heterozygous animals of lines #5 and #6 using real-time PCR. Expression of the transgene was observed in all tissues examined (brain, heart, intestine, kidney, liver, lung, testis). Fig. 2A shows the levels of transgene activity relative to the *Actb* gene in individual animal lines #5 and #6. In both lines the highest relative transgene activity was observed in the heart. One significant challenge is selecting a reference gene that is stable across different organs. Therefore, we interpret this result as a qualitative rather than quantitative finding that the transgene is active in all examined organs.However, it is important to emphasize that accurately measuring differences in gene activity between organs using real-time PCR is challenging.

**Figure 2.**
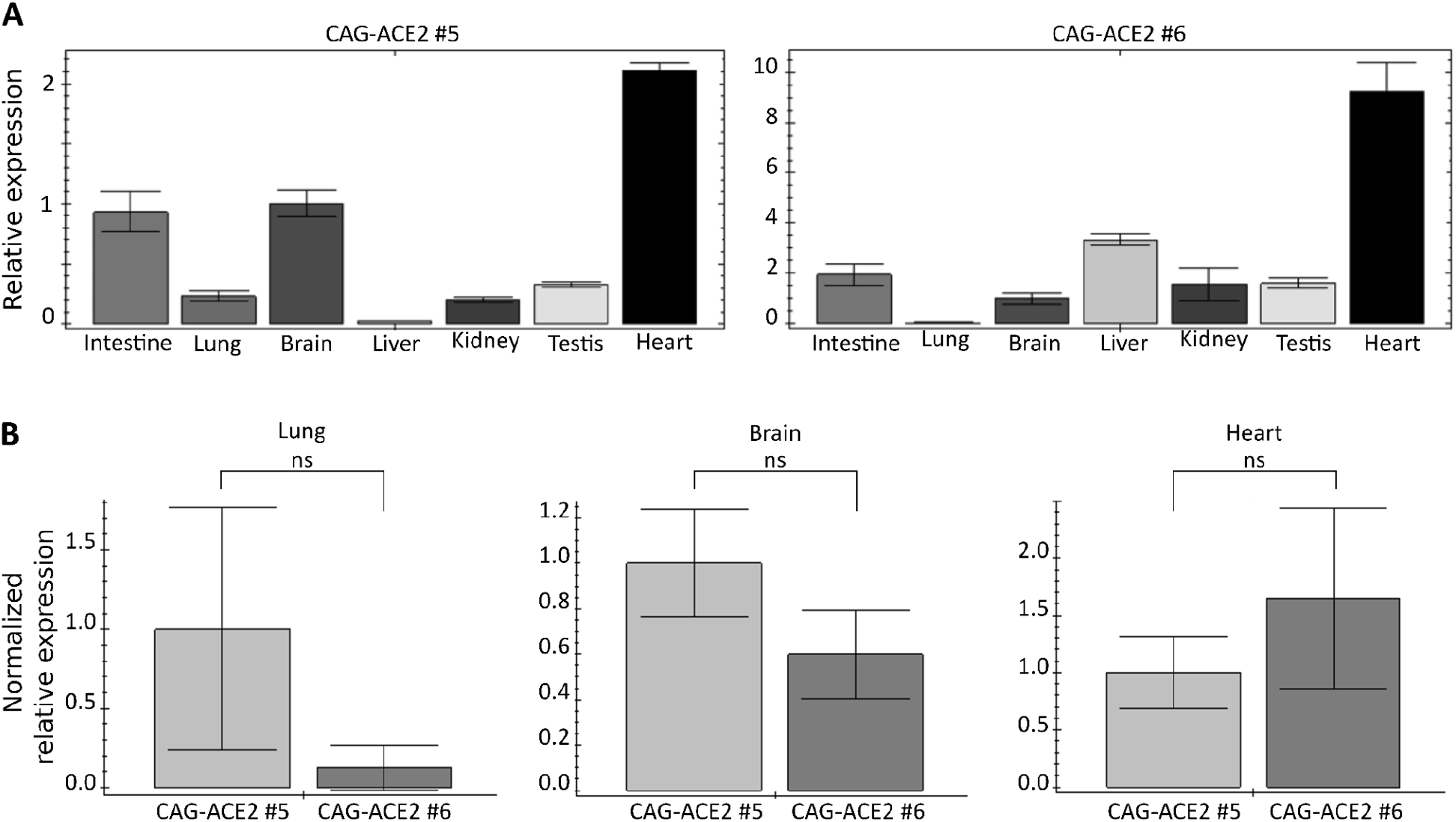
*Ace2* expression relative to *Actb* as a reference gene in various tissues of the two transgenic lines. (A) Expression profiles for the two transgenic lines. (B) Relative comparison for three tissue types between the lines. Sample values from #5 were set as 1. Data is shown as an average of three replicates and SEM.

To assess whether transgene activity differs between strains, we compared the average activity level of three animals of each strain in three organs: lung, brain, and heart (Fig. 2B). The analysis showed that there are no significant differences between the lines. Thus, the CAG promoter-based transgene provides ubiquitous activity of human *ACE2* in mouse organs.

### Transgenic mouse cells are susceptible to the SARS-CoV-2 virus

In order to evaluate whether transgene activity allows the SARS-CoV-2 virus to enter cells and support viral replication we infected primary cell cultures derived from mice. We isolated embryonic fibroblasts (MEFs) from E13.5 line #6 mouse embryos. We obtained two cell cultures from transgenic embryos and three cell cultures from non-transgenic siblings embryos.

Once the cells reached a subconfluent monolayer, infection was carried out with standard tenfold dilutions of a virus-containing suspension (infectious dose range 1000-0.01 of the 50% tissue culture infectious dose (TCID_50_)). After adsorption for 1 hour, the cells were washed and cultured for three days. Next, the presence and severity of virus-specific cytopathogenic effects were assessed qualitatively using light microscopy. In cell cultures of transgenic embryos infected with maximum multiplicity, clear signs of a virus-specific cytopathogenic effect were recorded 24 hours after infection, and after 72 hours with the entire range of infectious doses of SARS-CoV-2 used up to 0.1 TCID_50_. In other cell cultures obtained from non-transgenic embryos infected with any doses of SARS-CoV-2, no cytopathogenic effects were observed (Table 1).

**Table 1.** Infectious activity of the SARS-CoV-2 on MEF cells.

| Infectious dose of SARS-CoV-2, TCID<sub>50</sub> | MEFs from transgenic embryo 1 | MEFs from transgenic embryo 2 | MEFs from non transgenic embryo 1 | MEFs from non transgenic embryo 2 | MEFs from non transgenic embryo 3 |
| --- | --- | --- | --- | --- | --- |
|  | Severity of cytopathogenic effect (CPE) /virus titer (lgTCID<sub>50</sub>/ml*) |  |  |  |  |
| 1000 | ++++/6.4±0.4 | ++++/6.1±0.3 | ---/<1.0 | ---/<1.0 | ---/<1.0 |
| 100 | ++++/6.2±0.3 | ++++/6.3±0.4 | ---/<1.0 | ---/<1.0 | ---/<1.0 |
| 10 | ++++/6.5±0.3 | +++/6.4±0.4 | ---/<1.0 | ---/<1.0 | ---/<1.0 |
| 1 | ++/6.7±0.4 | ++/6.2±0.3 | ---/<1.0 | ---/<1.0 | ---/<1.0 |
| 0.1 | +/4.3±0.2 | +/3.9±0.3 | ---/<1.0 | ---/<1.0 | ---/<1.0 |
| 0.01 | ---/<1.0 | ---/<1.0 | ---/<1.0 | ---/<1.0 | ---/<1.0 |

(++++)- complete viral CPE (100% cell lysis); (+++) - 50-85% CPE; (++) - 20-50% CPE; (+) - single foci of viral CPE; (---) - complete preservation of the monolayer. \* - infectious titer of SARS-CoV-2 in the appropriate culture medium determined on Vero E6 cells (± SD for three independent experiments, p< 0.05).

To further confirm the specificity and evaluate the replicative activity of SARS-CoV-2 in the studied cell cultures, the culture media together with the cells were subjected to freezing, thawing and centrifugation, after which the infectious titers of SARS-CoV-2 were determined using a Vero E6 cells. As can be seen from the presented results (Table 1), SARS-CoV-2 is reproduced with high efficiency in cell cultures obtained from transgenic embryos, while no infectious SARS-CoV-2 was detected in cultures obtained from non-transgenic embryos.

In the Figure 3 monolayers of the transgenic MEFs were infected with various doses of SARS-CoV-2 illustrating that MEFs from transgenic embryos effectively support the replication of coronavirus and the ability of SARS-CoV-2 to cause a cytopathogenic effect with complete lysis of these cultures in wide range of infectious doses used.

**Figure 3.**
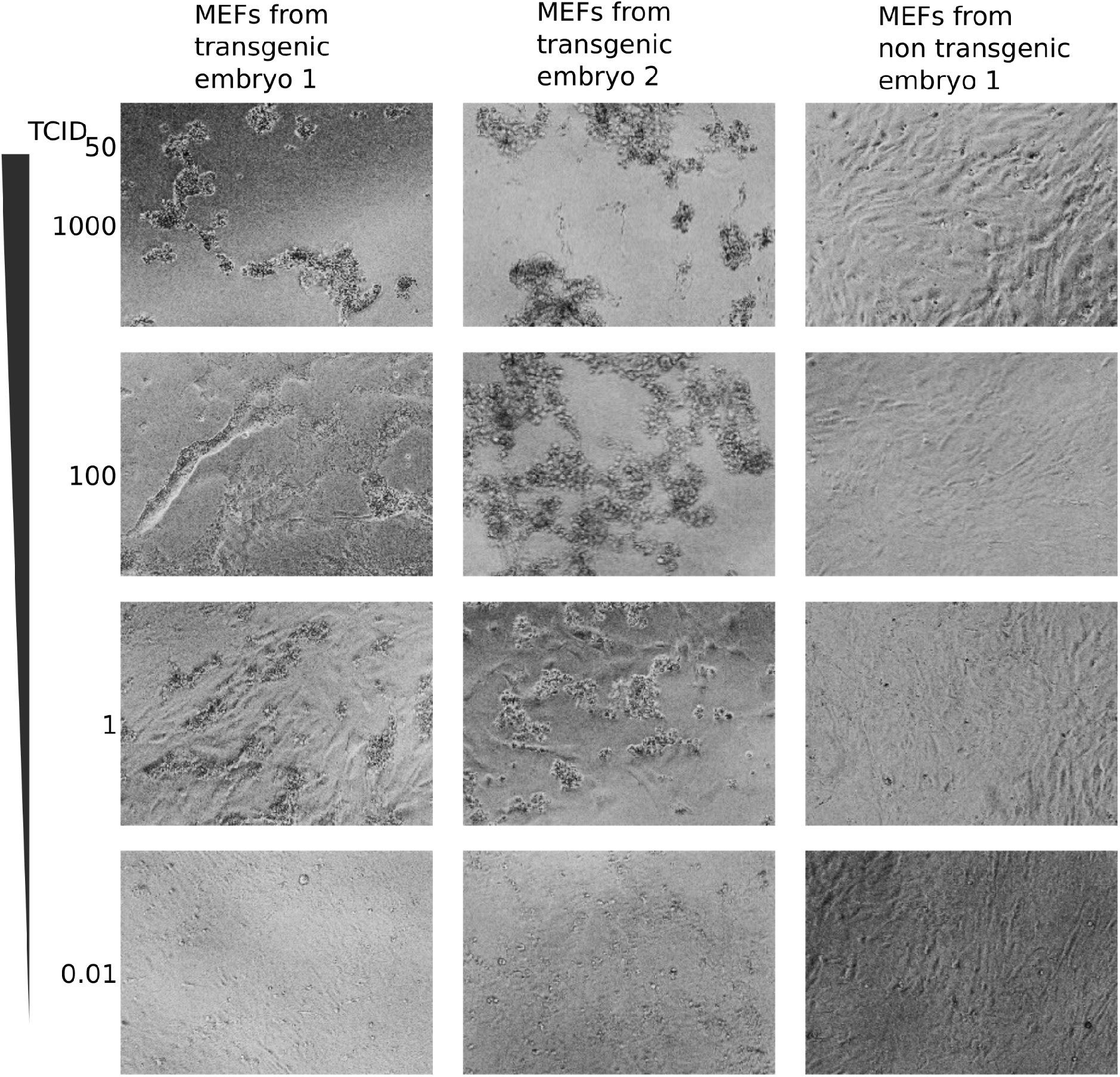
MEF cells from line #6 embryos are susceptible to infection. Representative views of MEFs cultures 72 hours after infection with the indicated virus load in TCID_50_.

Thus, the activity of the CAG-ACE2 transgene makes cells highly sensitive to lytic infection caused by SARS-CoV-2 and capable of effectively supporting coronavirus replication.

### Infection with SARS-CoV-2 leads to the death of transgenic CAG-ACE2 animals

In order to test the effect of SARS-CoV-2 on animals *in vivo*, we formed three experimental groups. Group I consisted of 10 transgenic females aged 8 weeks (5 females from line 5 and 5 females from line 6). This group of mice was intranasally infected with SARS-CoV-2 at a dose of 4.0 lg TCID_50_. Group II also consisted of 10 transgenic females aged 8 weeks (5 females of line 5 and 5 females of line 6). These animals were inoculated intranasally with PBS. Group III consisted of 6 non-transgenic wild-type females that were intranasally infected with SARS-CoV-2 at a dose of 4.0 lg TCID_50_.

No clinical manifestations were detected in the Group II and Group III. In Group I, starting from the third day after infection, there was a decrease in motor activity, hunching, rapid breathing, and a sharp loss of body weight; from day 4 – terminal condition, death. Last animals from Group I died after seven days of infection (Fig. 4). We collected the lungs and brains of 4 mice (2 just died and 2 in a terminal state) (Group I, day 5 after infection) to determine the presence of infectious SARS-CoV-2 and for pathomorphological examination. Organs from control mice (Group II and Group III) were also collected. First, we determined infectious activity of the SARS-CoV-2 in organ homogenates using Vero E6 cells. The SARS-CoV-2 virus was detected in a titer of 3.8±0.3 - 4.3±0.4 lg and 6.3±0.3 - 8.3±0.4 lgTCID_50_/g in the lungs and brains of 4 mice from Group I, respectively. In the lungs and brains of control animals (Group II and Group III), the infectious SARS-CoV-2 virus was not detected - lg TCID_50_/g - <0.5.

**Figure 4.**
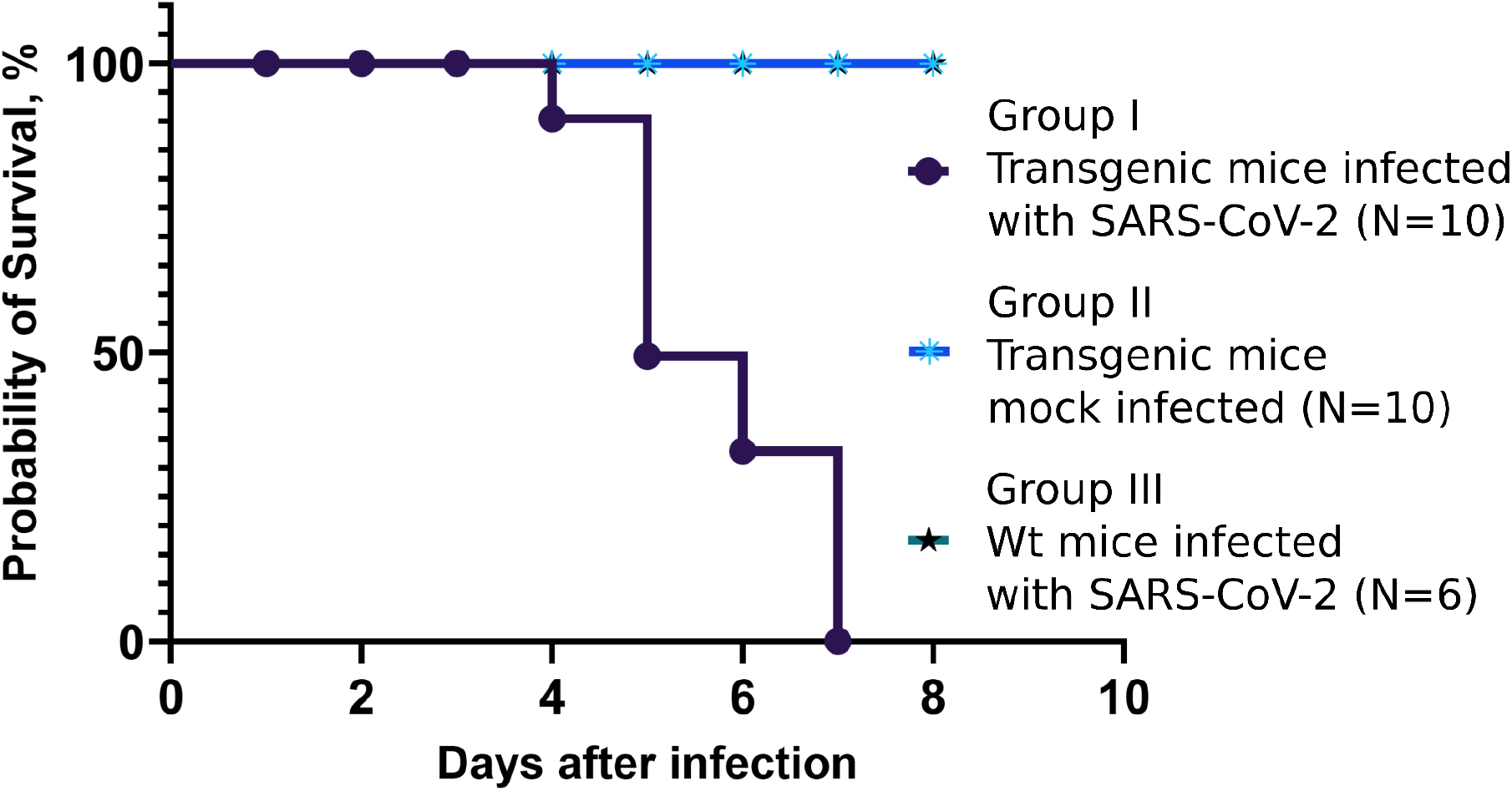
Transgenic animal survivability after SARS-CoV-2 infection. In groups Transgenic mice mock infected and Wt mice infected with SARS-CoV-2 there was not a single death of animals, so their graphs overlap.

A pathomorphological examination of mice from control animals (Group II and Group III) revealed no pathological changes in the lungs (Fig. 5 and data not shown). In mice of Group I, the following signs were detected in the lungs: a decrease in the airiness of the organ parenchyma (up to 50% of the section area); weakening of air filling is accompanied by a sharp plethora of blood vessels and large areas of hemorrhage; hemorrhages cover, upon visual assessment, from 40 to 50% of the parenchyma; inflammatory cellular reaction, represented by an increased number of neutrophils and lymphoid cells; focal dysplasia of the epithelium in the bronchi of medium caliber.

**Figure 5.**
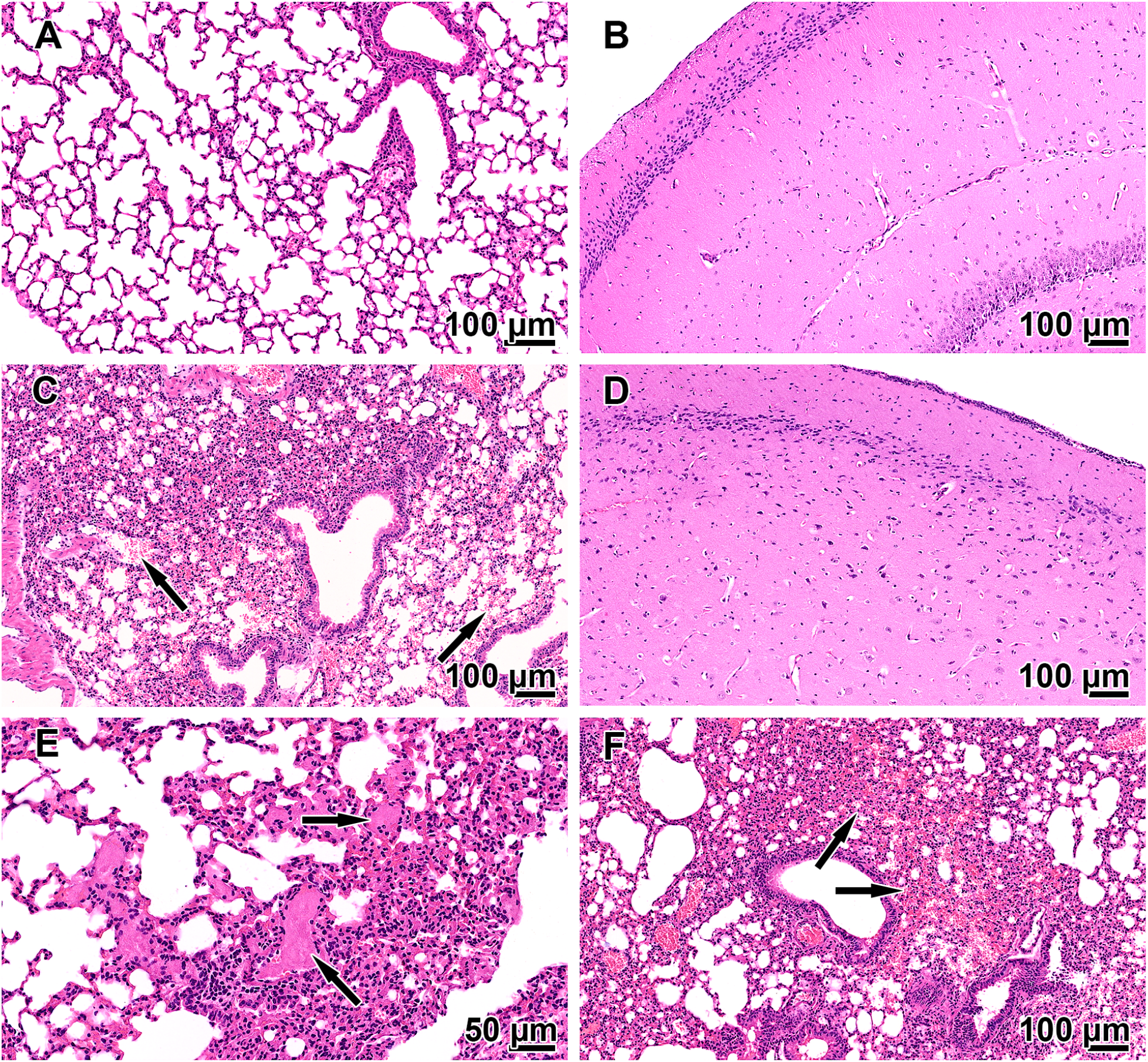
Pathological changes in the lungs and brains of the SARS-CoV-2 infected transgenic mice. (A) Group II (transgenic mock infected) normal lung morphology. (B) Group II (transgenic mock infected) normal structure of the cerebral cortex. (C) Group I (transgenic SARS-CoV-2 infected) decreased alveolar air volume, vascular hyperemia, blood cells in the lumens of the alveoli (arrows). (D) Group I (transgenic SARS-CoV-2 infected) normal structure of the cerebral cortex. (E) Group I (transgenic SARS-CoV-2 infected) decreased alveolar air volume, fibrin blood clots in blood vessels (arrows). (F) Group I (transgenic SARS-CoV-2 infected) vascular hyperemia and a large focus of intraalveolar hemorrhage (arrows).

In contrast to the lung samples, histological analysis of the brain did not reveal any significant changes among all groups of animals; only multiple sludge of erythrocytes was occasionally observed in the vessels of mice of the Group I (data not shown).

Thus, expression of the CAG-ACE2 transgene allows mice to be susceptible to the SARS-CoV-2 virus and thus provides a lethal animal model of COVID-19.

### Analysis of the concatemer structure in mice line #6

In addition to the functional elements of the transgenic construct, the structure of the transgenic locus itself can influence the properties of transgenic animals. It is known that during pronuclear microinjection, the introduced copies of linear DNA are united in tandem (Smirnov et al. 2019). We assessed the copy number of the transgenic construct among the line founders using the Droplet Digital PCR (ddPCR) method. Line #5 had 140 ± 27 copies in the genome, #6 - 72 ± 8, #10 - 73 ± 7. Inside the concatemer most copies are usually oriented in a head-to-tail direction, but palindromic junctions and truncated transgene copies could also be present (Smirnov and Battulin 2021). However, due to its repeated nature, the internal concatemer structure is extremely difficult to study. With the advent of third-generation sequencing methods (Oxford Nanopore and PacBio) capable of reading individual DNA molecules tens and even hundreds of thousands of base pairs in length, we have become much better able to understand the actual structure of transgenic loci.

In order to characterize the structure of the transgene in the CAG-ACE2 line #6, we used sequencing on the Oxford Nanopore platform. However, it is worth emphasizing that even in the case of a high copy number of a concatemer, it constitutes only a small part of the entire genome. So for line #6 the length of the concatemer will be about 360 kb (5 kb * 72 copies), which is only 0.012% of the haploid mouse genome. Accordingly, when sequencing the genome of a transgenic animal, only 0.012% of the reads will carry information about the DNA sequence in the transgenic locus. We performed genomic sequencing of a DNA sample isolated from a F1 heterozygous carrier from line #6 using Oxford Nanopore with ∼0.25x coverage. We obtained 15 reads mapping to the CAG-ACE2 plasmid (Fig. 6A). Among these reads, the most informative was a ∼40 kb long read consisting entirely of concatemer sequence. The dot-plot alignment of this read to the plasmid sequence is presented in Figure 6B. It is clearly seen that the read covers 7 full transgene copies and one truncated copy of the transgene that has lost the end of the ACE2 coding part and the polyadenylation signal. The copies are arranged in a head-to-tail orientation. In this read, the backbone fragment was also discovered, which probably co-purified with transgene DNA during gel extraction (Fig. 6B). Although this long read gives a good idea of the structure of the concatemer, there is insufficient data for quantitative characterization. Therefore, we applied nCATS Cas9 single cut approach method in order to increase the proportion of informative reads covering the transgenic locus (Gilpatrick et al. 2020). We isolated high-molecular weight DNA from line #6 spleen, dephosphorylated the ends, and cut with Cas9 *in vitro* inside the middle of the transgene (Fig. 6C). When further preparing libraries for sequencing, adapters for nanopore sequencing are preferentially ligated to the ends formed after Cas9 cleavage, since these ends have phosphates. This allowed us to repeatedly enrich the data with reads with a transgene insertion, in our case more than 100 times. For further analysis, we selected only reads that map to the transgene and are longer than 3000 bp. This constituted 864 reads. Inspection of these reads revealed the internal structure of the transgenic locus in detail. In particular, we found fragments in the reads obtained by the Cas9 single cut enrichment method corresponding to sections of a long 40 kb read obtained by whole-genome nanopore sequencing (Fig. 6B, C). The large amount of obtained data made it possible to quantify the internal junction structure of the transgenic locus. In sum, we found 1521 segments (segment = transgene copy or plasmid backbone fragment) (over 200 bp in length) of the concatemer, 1373 copies from the target transgene, and 148 from the excised plasmid backbone. Thus, about 10% of the concatemer segments originate from the plasmid backbone. The vast majority of transgene copies were joined in a head-to-tail orientation. However, we found 23 reads in which there was a palindromic connection of the ends of the concatemer copies. This is about 1.5% of all junctions between segments (Fig. 6D). Finally, we found a transgene genomic boundary: 6 independent reads show that the transgene was integrated into a region of the first chromosome. In this case, the flanking copy of the transgene was not truncated (Fig. 6E). Interestingly, the second concatemer border was not found, probably because it was close to a Cas9 cut site and can’t be attributed to transgene reads.

**Figure 6.**
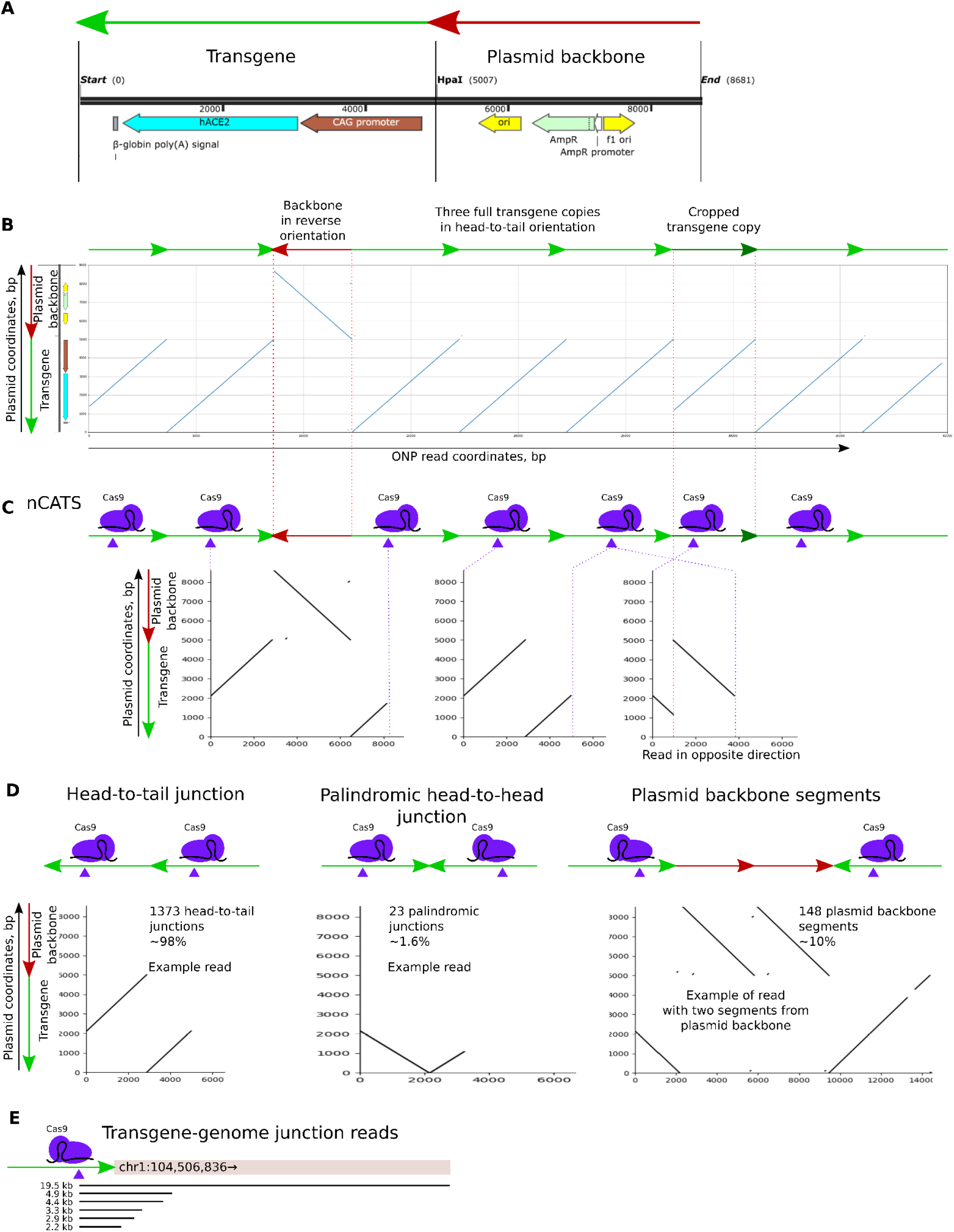
Transgene structure. (A) Basic elements of the plasmid into which the transgene was cloned. (B) Dot-plot alignment of a long nanopore read (x-axis) onto a plasmid (y-axis). The read covers a region of the concatemer containing 8 copies of the target transgene. (C) Fragments from the concatemer in section B cut by Cas9 for nCATS library preparation, along with the resulting reads. (D) Characteristic structures found in nCATS reads. (E) Reads covering the transgene-genomic junction found in nCATS reads.

## Discussion

Here we described the establishment of a lethal mouse model susceptible to SARS-CoV-2. Currently, several transgenic mouse lines have been created to model COVID-19. By far the most common model was the CK18-ACE2 line, created before the start of the pandemic. That line is great for modeling a wide variety of aspects of COVID-19. However, problems with the availability of these animals during the pandemic have prompted many groups to begin creating independent transgenic models. We believe that conducting studies in a variety of independently derived animal models provides more reliable data for the development of this field of science. Since in this case it is possible to separate the effects caused by the individual characteristics of the model from the really important effects inherent in the modeled process.

Through experimental infections with SARS-CoV-2 *in vitro*, we found that MEFs from transgenic mouse embryos are able to support coronavirus lytic infection with an efficiency not inferior to the highly sensitive to SARS-CoV-2 of Vero E6 cells. Here, using CAG-ACE2 transgenic mice, we show that infection with SARS-CoV-2 causes severe lethal disease. CAG-ACE2 mice infected with SARS-CoV-2 developed disease with features (clinical manifestations, level of mortality, SARS-CoV-2 replication activity in lungs and brains, tissue injuries) which are similar to that described earlier for CK18-hACE2 (Zheng et al. 2021) or UBC-ACE2 (Dolskiy et al. 2021) transgenic mice.

We decided to use the ubiquitous constitutive CAG promoter to express the human coronavirus receptor ACE2 in the widest possible range of cell types. The analysis of a set of animal organs showed the activity of the transgene in all organs studied. It is worth noting that the level of ACE2 expression differs among the organs, with the maximum level of expression observed in the heart. A similar pattern of activity with maximum transgene expression in skeletal muscle and heart was also previously noted in a line of transgenic mice with Carbonyl reductase 1 driven by the CAG promoter, so this may be a property of the promoter (Yokoyama et al. 2023). It is worth noting that, as far as we know, there is no information yet on how the level of ACE2 expression affects the course of the disease in animals, and the threshold level of expression that will ensure the susceptibility of animals to the virus is also not known. Of course, such information would be extremely useful for the rational design of a transgenic model.

CAG has proven to be a successful choice for driving widespread constitutive transgene expression in animals. Recently, work was published with a similar design in which CAG drove ACE2 expression, and the resulting animals, like those described in this article, are a lethal model of COVID-19 (Liu et al. 2024). It is worth emphasizing that the choice of promoter greatly influences the properties of the construct. So, we previously tried to make a similar model using the EF1a promoter, but all lines of transgenic animals obtained in this way turned out to be immune to SARS-CoV-2 due to silencing of the construct in animal tissues (Battulin et al. 2022). The peculiarities of the EF1a and CAG promoter were well demonstrated in the work on creating humanized pigs for xenotransplantation purposes (Anand et al. 2023). A cassette containing both EF1a and CAG promoters is integrated into the genome of these animals. Nanopore direct RNA-seq of kidney mRNA in that article shows that there are a hundred times fewer transcripts with the EF1a promoter than transcripts with the CAG promoter. Importantly, the difference appears precisely *in vivo* during the development of a living organism, since when compared on cells, both of these promoters are among the most powerful, in cells of different tissue origin. This once again emphasizes the importance of the correct choice of promoter for creating transgenic animals.

To characterize the transgenic locus, we used sequencing on the Oxford Nanopore platform with Cas9 single cut enrichment. It is known that as a result of pronuclear microinjection of linear DNA, a tandem of several copies of the introduced DNA is usually inserted. We have previously shown that such association occurs due to the mechanism of homologous recombination. The number of transgene copies in the concatemer and the integration site can affect the activity of the construct, but there is not much systematic data on the structure of the transgene locus. The main difficulty is the repeated nature of the concatemer. Since the typical size of an NGS read is usually tens of times smaller than the characteristic size of a transgene, it is impossible to determine the structure of a repeat consisting of transgenes using NGS. Progress in this area became possible with the advent of third-generation sequencing methods, in which the read length is greater than the size of one copy of the concatemer (Jupe et al. 2019).

In our case, in line #6, about 70 copies of the transgene (according to ddPCR data) were inserted into the intergenic region of mouse chromosome 1 (Fig. 6E). Most copies of the transgene are oriented in a head-to-tail configuration. To our knowledge, the molecular mechanism leading to such asymmetry has not yet been described; this is one of the fundamental unsolved mysteries in transgenesis (Smirnov and Battulin 2021). Hypothetically, homologous recombination can direct ligation of the transgenes’ ends into head-to-tail orientation. Another possibility is that all copies are ligated randomly by non-homologous or microhomology-mediated end-joining (NHEJ/MMEJ) but palindromic transgene junction variants (head-to-head and tail-to-tail) can form toxic secondary structures in genomic DNA such as hairpins or cruciforms. Apparently, long palindromes are unstable in the genome; for example, the occurrence of deletions that destroy the point of symmetry of a palindrome, which was formed from two copies of a transgene oriented head to head, has been described. In our data, we also observed this copy orientation in 1.6% of sequenced internal junctions. The stability of these structures in our case is not yet known.

We also found that between some copies of the target transgene in the concatemer there was integration of plasmid sequences that apparently could not be eliminated 100% when purified from an agarose gel. Fragments of the bacterial genome and plasmid backbone that are present in the injection mixture can be incorporated into the concatemer, for example, a fragment of the E. coli genome previously was found in the concatemer of the Oct4-EGFP transgene (Nicholls et al. 2019). It can be assumed that such off-target insertions of bacterial DNA are quite common among transgenic animals, but they are not easy to detect unless specifically studied.

We found only a few copies of the transgene with terminal deletions, which indicates a high integrity of the introduced constructs. Truncated copies also appear to be quite common among transgenic animals; for example, such copies were described among 29 copies of the Oct4-EGFP transgene (Nicholls et al. 2019). And K18-ACE2, according to nanopore sequencing of 10 complete concatemers, contains 10 complete copies of the transgene and 4 with various large deletions (Low et al. 2022).

Overall, the transgene-cutting Nanopore enrichment approach turned out to be very informative. It can be recommended for describing the internal structure of the transgene.

In conclusion, we have successfully generated a murine model that expresses human ACE2 through a ubiquitous CAG promoter, making it highly susceptible to SARS-CoV-2 infection. This model demonstrated rapid and lethal disease progression, mirroring severe human COVID-19, making it a valuable tool for studying the virus and testing potential treatments. The transgene was active in all examined tissues, and supported effective viral replication. Additionally, we characterized the structure of the transgene locus using advanced sequencing techniques, revealing a high-copy concatemer primarily in a head-to-tail orientation. This model, alongside others, will enhance our understanding and facilitate the development of therapeutic strategies against COVID-19.

## Materials and methods

### Transgenic mice generation

The CAG-ACE2 genetic construct was prepared on the basis of the pT2/BH backbone (BglII, EcoRI sites) (Addgene #26556). The promoter region from the pCAGGS-mCherry vector (Addgene #41583) was combined with the human ACE2 CDS (Addgene #1786) and the three polyadenylation signals (b-globin, SV40, bGH). Prior to microinjection, plasmid DNA was treated with HpaI to excise the ACE2 gene with the CAG-promoter and the b-globin polyA signal (total length 5007 bp) (Fig. 1A). DNA was gel purified and diluted to 6 ng/μL in the TE buffer. The solution was backfilled into an injection needle with positive balancing pressure (Transjector 5246, Eppendorf) and injected into the cytoplasm of zygotes of C57BL/6 background. After injections, the embryos were cultured for 1 h in drops of M16 medium at 37°C and an atmosphere of 5% CO2. The viable microinjected zygotes were transplanted the same day into oviducts of pseudopregnant CD-1 females (0.5 days after coitus). Isoflurane inhalation anesthesia was applied in these experiments.

Transgene insertion in mice was confirmed by PCR genotyping with primers for ACE2 gene (Table 1). For ddPCR, primers and probes for ACE2 and mouse Emid1 reference genes (one copy in mouse haploid genome) (Table 1) were used in accordance with manufacturer’s protocol (ddPCR Supermix for Probes (No dUTP), BioRad). Droplet digital PCR (ddPCR) was performed using a QX100 system (Bio-Rad). In brief, genomic DNA was digested overnight with MseI in CutSmart buffer (NEB) (1 μg genomic DNA in 30 μl) and added to the ddPCR mixture. ddPCR reactions were set in 20 μl volumes containing 1× ddPCR Supermix for Probes (no dUTP), 900 nM primers and 250 nM probes, and 1 μl of genomic DNA. Amount of DNA was based on the copy number for the exact line, and was in the range of 0.3-30 ng per reaction. ddPCR reactions for each sample were performed in duplicates. PCR was conducted according to the following program: 95 °C for 10 min, then 41 cycles of 95 °C for 30 s and 61 °C for 1 min, with a ramp rate of 2 °C per second, and a final step at 98 °C for 5 min. The results were analyzed using QuantaSoft software (Bio-Rad).

### Viruses and cell cultures

In early 2020, the hCoV-19/Russia/StPetersburg-64304/2020 (GISAID, EPI_ISL_428868) SARS-CoV-2 strain was isolated from Vero E6 cells in Russia from a patient suffering from COVID-19 (Svyatchenko et al. 2021).

Vero E6 cells were obtained from the SRC VB “Vector” (Russia) collection and were grown using DMEM (BioloT Ltd., Russia) in the presence of 10% fetal bovine serum (HyClone, USA), penicillin (100 IU/mL), and streptomycin (100 μg/mL).

The viral titers for SARS-CoV-2 in Vero E6 cells were determined using a 50% tissue culture infectious dose (TCID_50_) assay as estimated by microscopic scoring of the cytopathic effect and by measuring cell viability in the formazan-based MTT assay as described previously (Niks and Otto 1990). Viral stocks of SARS-CoV-2 with infectious titers of 6.5 lg TCID_50_/ml were stored at −70 °C.

### Animals

The experimental animals were fed a standard diet and had access to water *ad libitum*, according to veterinary legislation and the requirements for humane animal care and the use of laboratory animals (National Research Council, 2011). The animal experiments were approved by the Bioethics Committee of the State Research Center of Virology and Biotechnology “Vector”.

### Infection with SARS-CoV-2 *in vitro*

MEF cells were cultured in 24-well plates using DMEM (BioloT Ltd., Russia) in the presence of 10% fetal bovine serum (HyClone, USA), penicillin (100 IU/mL), and streptomycin (100 μg/mL) and inoculated in triplicate with SARS-CoV-2 in different dilutions. The SARS-CoV-2 infectious viral titers were estimated by microscopic scoring of the CPE and by measuring cell viability in the formazan-based MTT assay.

### Infection in vivo

SARS-CoV-2 infection was studied *in vivo* using transgenic mice. The animals were intranasally challenged with SARS-CoV-2 (10^4^ TCID_50_) and mock-infected animals were used as control. Intranasal inoculation was performed by anaesthetizing the mice with isoflurane (Isothesia; Henry Schein Animal Health) and then inoculating the nostrils with the viruses in 50 µL of phosphate buffered saline (PBS). The control animals were administered with PBS. After the challenge, the animals were observed and weighed daily.

On day 5 post infection, the brain and lungs were collected from experimental animals. The infected tissues were homogenized and used to determine the infectious titers of the virus. The SARS-CoV-2 titers, expressed as the 50% tissue culture infectious doses (TCID_50_), were determined by the cytopathic effect (CPE) assay in Vero E6 cells. The tissues from experimental mice were also fixed in formalin and used for the pathohistological study (Svyatchenko et al. 2023). The slide samples were stained using a standard haematoxylin–eosin staining procedure. Light-optical examination and microphotography were carried out using an Imager Z1 microscope (Zeiss, Göttingen, Germany) equipped with a high-resolution HRc camera. The images were analyzed using the AxioVision Rel.4.8.2 software package (Carl Zeiss MicroImaging GmbH, Jena, Germany).

Animal euthanasia was carried out using an automated compact CO2 system for humane output from the experiment of laboratory animals (Euthanizer, Russia). The concentration of carbon dioxide (30% at the 1st stage, 70% at the 2nd stage) and gas supply rate satisfy the requirements of the American Veterinary Medical Association (AVMA) 2020.

### Statistical analysis

Basic statistical analyses, including calculations of the mean, standard deviation, and coefficient of variation of the mean Ct value, were performed using Excel (Microsoft Corp., Redmond, WA, USA). Statistical data processing was conducted using the statistical program STATISTICA 12 (StatSoft Inc., USA), also Graphpad Prism (GraphPad Software, Inc., USA) was used to analyze expression data. Statistical evaluation of the differences between the groups was performed using the Student’s t-test and the Mann–Whitney U test for expression data. p < 0.05 was considered significant.

### Biosafety

All experiments involving any infectious viral materials were conducted in a Biosafety Level-3 Laboratory with all applicable national certificates and permissions required for studying SARS-CoV-2.

Institutional Review Board Statement: The animal study was conducted according to the guide-lines of the Declaration of Helsinki, in compliance with the protocols and recommendations for the proper use and care of laboratory animals (ECC Directive 86/609/EEC, Ministry of Health of Russian Federation Directive no. 708n/2010, and Ministry of Health of Russian Federation Directive no. 267/2003). The protocol was approved by the Bioethics Committee of the Federal Budgetary Research Institution State Research Center of Virology and Biotechnology ”Vector” (protocol code: SRC VB “Vector”/02-05; date of approval: 05.15.2020).

### Isolation of total RNA and analysis of ACE2 expression by qPCR

Tissue samples were isolated from mice and stored in a -80°C freezer. The following organs were used to isolate total RNA: brain, heart, lung, liver, intestine, kidney, testis. RNA was isolated using QIAzol reagent (QIAGEN Science, USA) according to the manufacturer’s protocol. To synthesize the first strand of cDNA, 2 μg of isolated RNA, previously purified from genomic DNA using DNAse l (Thermo Fisher Scientific, USA), was selected. 1 μg of purified RNA was taken into a new tube for further use as an RT-control. Synthesis of the first strand of DNA was carried out using the “Reverse transcriptase M-MuLV-RH” kit (Biolabmix, Russia) and random hexaprimers in accordance with the manufacturer’s protocol. 1 μl of cDNA was used as a matrix, as well as the following primers and probes for ACE2 and ActB as a reference gene (Table 2). qPCR was performed using the BioMaster HS-qPCR(2x) kit (Biolabmix, Russia). For each sample with its own set of primers and probes, 3 replicates were carried out. The reaction was carried out on a Real-time CFX96 Touch thermal cycler (Bio-Rad, USA) under the following conditions: Pre-denaturation 95°C for 5 minutes, then 39 cycles of denaturation 95°C 10 seconds, primer annealing 60°C 30 seconds, elongation 72°C 30 seconds. Using CFX Manager software (Bio-Rad, USA), graphs of relative normalized expression (ΔΔCq) were obtained for each sample.

**Table 2.**
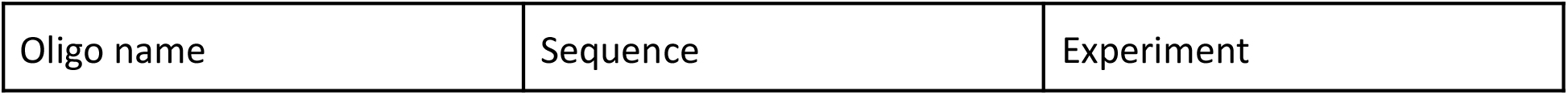
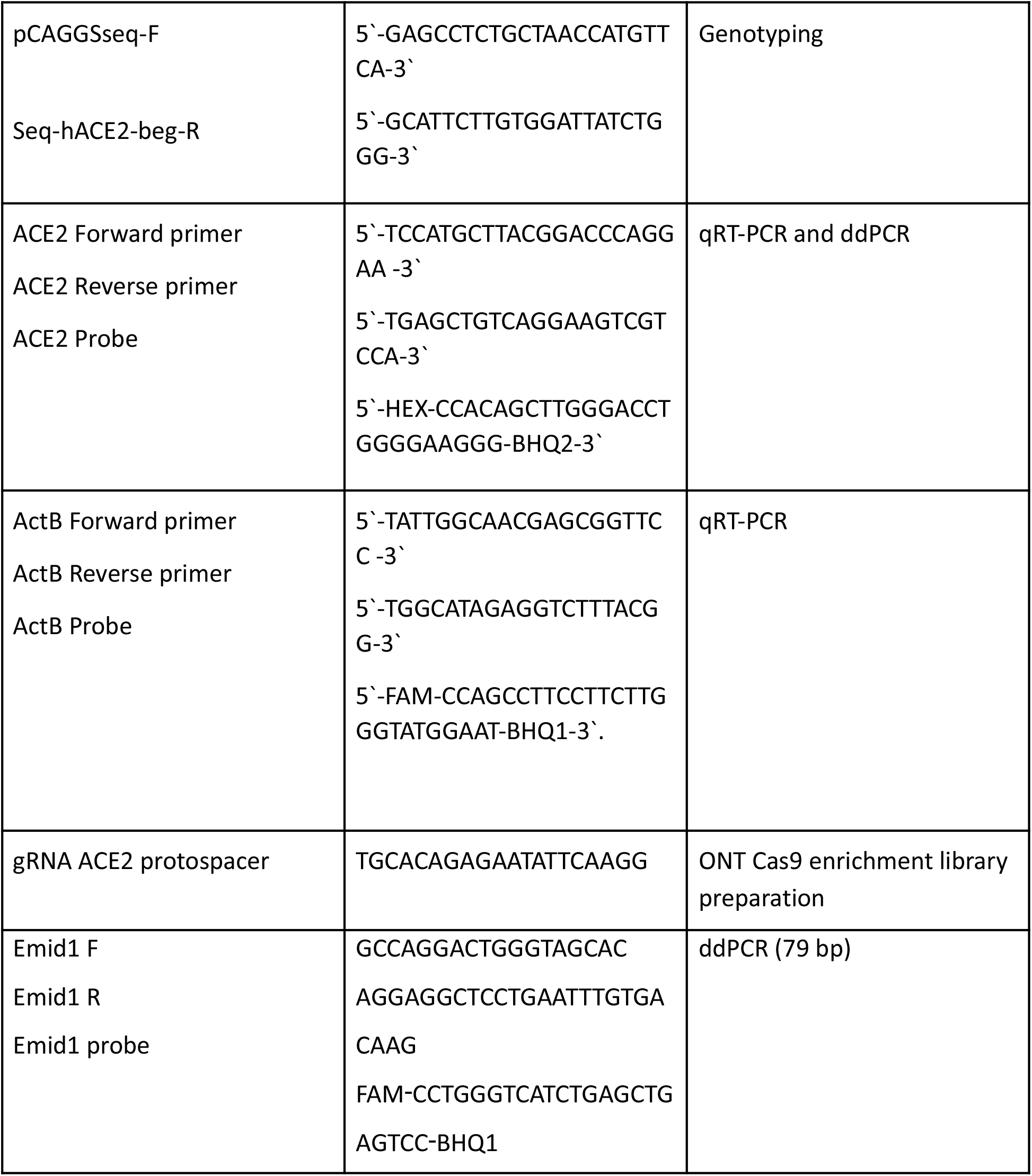
Primers and sgRNA sequences.

### ONT Cas9 enrichment library preparation

High-molecular weight DNA was extracted from mouse tissues as described in (Quick 2018) and homogenized by passing through G26 needle 10-15 times. Extracted DNA was left for dissolving at 4°C overnight. Library was prepared according to standard “Ligation sequencing gDNA - Cas9 enrichment” protocol for SQK-LSK109 kit with minor changes in enzymes and buffers as products from alternative manufacturers were used. sgRNA complementary to the central part of the hACE2 (Table 2) was synthesized with HiScribe™ T7 High Yield RNA Synthesis Kit (NEB, USA). Library was loaded onto the FLO-MIN106 flow cell and sequenced with GridION (Oxford Nanopore, UK).

## Author Contributions

Conceptualization, N.B., A.S., V.B.L. and O.S. ; methodology, A.S., A.N., G.K., I.S., E.K., E.C., E.M., M.S., D.Z., V.A.S., E.V.P., O.S.T., S.S.L.,; writing—original draft preparation, N.B.; writing—review and editing, N.B., A.S. and A.N.; supervision, N.B. and O.S.; funding acquisition, N.B. and O.S. All authors have read and agreed to the published version of the manuscript.

## Acknowledgement

We thank the Talent and Success Foundation for supporting the work on characterizing the structure of the transgene using the ONT sequencing method, which was carried out during the “Grand Challenges” schoolchildren scientific and technological project programme. Specifically, we thank Angelina Veklenko, Irina Karaman, Daria Perfilyeva, Veronika Reznichenko, Ayana Khankharaeva, Matvey Shavernev - schoolchildren who contributed to this work.

